# Metabolic modelling reveals increased autonomy and antagonism in type 2 diabetic gut microbiota

**DOI:** 10.1101/2024.07.31.605966

**Authors:** A. Samer Kadibalban, Axel Künstner, Torsten Schröder, Julius Zauleck, Oliver Witt, Georgios Marinos, Christoph Kaleta

**Author notes:** Senior authorship; to whom correspondence should be addressed.

## Abstract

Type 2 diabetes presents a growing global health concern, with emerging evidence highlighting the pivotal role of the human gut microbiome in metabolic diseases. This study employs metabolic modelling to elucidate changes in host-microbiome interactions in type 2 diabetes. Glucose levels, dietary intake, 16S sequences and metadata were estimated and collected for a cohort of 1,866 individuals. In addition, microbial community models, as well as ecological interactions were simulated for the gut microbiomes of the cohort participants. Our findings revealed a significant decrease in the fluxes of metabolites provided by the host to the microbiome through the diet in patients with type 2 diabetes, accompanied by an increase in within-community exchanges. Moreover, the diabetic microbial community shifts towards increased exploitative ecological interactions among its member species at the expense of collaborative interactions. The reduced butyrate flux from the community to the host and reduced tryptophan acquired by the microbiome from the host’s diet further highlight the dysregulation in microbial-host interactions in diabetes. Additionally, microbiomes of type 2 diabetes patients exhibit enrichment in energy metabolism pathways, indicative of increased metabolic activity and antagonism. This study provides insights into the metabolic dynamics of the diabetic gut microbiome, shedding light on its increased autonomy and altered ecological interactions accompanying diabetes, and provides candidate metabolic targets for intervention studies and experimental validations, such as butyrate, tryptophan, H2S, several nucleotides, amino acids, and B vitamins.

## 2. Introduction

Despite the increased awareness as well as the development and availability of diagnostic and treatment approaches, type 2 diabetes (T2D) still presents an increasing global health issue. It was estimated that in 2021, around 537 million adults were living with diabetes, and this is forecasted to reach 783 million by 2045 (1). The global increase in T2D is facilitated by the continuous transition into an industrialised lifestyle that intrinsically affects people’s daily activities and dietary habits. The human gut is one of the densest known microbial habitats. With a high biodiversity and a genetic repertoire of over three million genes and thousands of metabolites (2), such an ecosystem is characterised by a complex network of metabolic interactions between the microbial community and the host. This crosstalk shapes the relations among the microbial species and plays a crucial role in the host’s health.

Moreover, the gut microbial community has a profound influence on human metabolism, which becomes particularly important in the context of metabolic diseases such as diabetes, where changes in the gut microbiota are associated with altered glucose homeostasis, which is intricately linked to the development of type 2 diabetes and its related complications (3). In addition, the impact of the gut microbiota extends to various physiological processes, including chronic low-grade inflammation, β-cell dysfunction, gut permeability, glycolipid metabolism, and insulin sensitivity, all of which are pertinent to T2D development (4). The mutual influence between T2D and the gut microbial community is bidirectional, as elevated blood sugar levels can heighten intestinal permeability via a glucose transporter, altering the gut microbiome and triggering an inflammatory response (5). Moreover, there’s a growing body of evidence suggesting that it’s not only the microorganisms themselves but also their metabolites that contribute to the development of metabolic syndrome and diabetes (6). Hence, understanding the complex relationship between the gut microbial community and diabetes could potentially lead to new strategies for prevention and treatment. However, while certain bacterial genera, such as *Ruminococcus*, *Fusobacterium*, *Blautia* and *Clostridium* have been reported in higher abundances among patients with T2D (3,7), more research is needed to fully understand the complex interactions and interplay between pathomechanisms of T2D and the microbial community. Moreover, the influence of host-derived metabolites on the gut microbial community still needs to be more thoroughly investigated (8).

Genome-scale metabolic models are a powerful tool that provides a comprehensive representation of the entire metabolic network of an organism. Those models are reconstructed by associating the organism’s genomic content with enzymatic reactions and metabolic transport (9). The use of metabolic modelling through flux balance analysis is being increasingly employed to investigate complex metabolic networks and to study the metabolic interactions within microbial communities and with the host, especially in complex diseases such as Parkinson’s and Alzheimer’s (10,11) as well as IBD (12,13). Nonetheless, the use of metabolic modelling to understand the alterations in the metabolic exchange profile between the gut microbiome and the human host in the context of diabetes is still limited to a few attempts (14–16). Arguably, glucose stands out as the most crucial energy carrier and carbon source, serving as a foundation for metabolites and the construction of biopolymers across all kingdoms of life

(17). In this study, the gut microbiome differential species abundances for a cohort of diabetic and healthy individuals were investigated. Thereafter, metabolic modelling was employed to predict the ecological interactions among the gut microbiomes as well as the microbial community metabolic fluxes. Subsequently, the microbiome metabolites associated with blood glucose measures and the production of those metabolites by dominant bacterial genera were observed. Finally, we hypothesise that there is an increased availability of glucose in the gut environment accompanying diabetes (18). The high availability of glucose as a simple energy source for the bacteria in the gut alters the microbial ecological interactions and metabolic exchanges. Those alterations include an increase in antagonism, energy metabolism and autonomy of the gut microbiome, which becomes less shaped by the host’s diet and more dependent on metabolic exchanges among its species.

## 3. Methods

### 3.1. Collection of metadata and 16S sequences

#### Metadata

In this study, Perfood established a cohort dataset comprising 2,117 individuals who purchased a Perfood product and provided consent for the anonymous use of their data for research purposes. Following the exclusion of cancer patients, individuals with inflammatory bowel disease (IBD), those undergoing antibiotic treatment during sample collection and those with repeated measures, the final dataset consisted of 1,866 individuals, with each participant contributing a single sample.

The cohort comprised 81 patients with type 2 diabetes and 1,785 non-diabetic individuals. Clinical metadata, including gender, age, BMI, waist-to-hip ratio, physical activity, medication against diabetes and other diseases and clinical measures, were collected using questionnaires (Supplementary Figure S1). Physical activity was estimated by aggregating the daily duration across all types of physical activities the participants undertook. Baseline blood glucose levels (before meals) were estimated using a pipeline developed in-house by Perfood, patented and published (19), and the estimated glucose levels were curated manually thereafter. In addition, HbA1c levels were collected using continuous glucose monitoring (CGM) sensors and averaged for the aim of this study.

#### Sequence data

Faecal samples were obtained from each participant and processed as described by Kordowski and others (20). The 16S rRNA gene sequences were acquired using Illumina paired-end short-read sequencing. Prior to downstream analysis, adapter sequences were trimmed, and the 16S sequences were filtered for host sequence contamination and screened for errors, where regions with high variability in k-mer frequencies, indicative of potential sequencing errors, were identified. The majority of the 16S sequences exhibited low error rates, less than 0.05, and sequences with higher error rates were excluded. Subsequently, the forward and reverse sequences were merged. These steps were performed using the R package dada2 (21). A list of all functions utilised is available in supplementary table S1.

#### Precision diet

Dietary information for each participant was collected through the Perfood digital diary platform. The total protein, carbohydrate, fats, and caloric intake from these dietary entries were estimated using an in-house dietary ingredient database. A reference precision diet, curated and adjusted to ModelSEED namespace (22) (referred to as the FOCUS diet), was used to estimate the expected frequencies of each macro– and micro-nutrient for the participants in the study. The process involved determining the proportion of each amino acid, fatty acid, sugar, and micro-nutrient (vitamins or minerals) based on the total protein, fat, carbohydrate, and caloric content within the FOCUS cohort. These proportions were then averaged across participants of the FOCUS cohort to establish reference ratios. Subsequently, these averaged ratios were applied to the Perfood cohort to estimate the frequencies of micro– and macro-nutrients for each participant. This estimation was achieved by multiplying the reference dietary element proportions by the estimated total protein, carbohydrate, fats, and caloric intake within their respective categories. The result is an estimated precision diet, ready for integration into community models after normalisation by caloric intake.

### 3.2. Microbial diversity and ecological interactions

The 16S sequences were mapped against a reference collection of the human gut microbiome (AGORA) (23) using the USEARCH software (24). We used the whole genomes of bacteria from the AGORA collection identified by the mapping procedure to construct genome-scale metabolic models using gapseq (9). Upon mapping the 16S sequences against the AGORA reference collection, we calculated the abundances of individual bacterial species within each microbial community. This was achieved by counting the 16S sequences that mapped to a certain genome for each participant and dividing this number by the sum of all 16S sequence counts within the community. We thereafter removed bacterial species with low abundances (below 0.1%) from the communities. This resulted in 499 bacterial species belonging to 148 genera that are present in different abundances within the gut microbial communities of the participants. Three alpha diversity measures, namely, the Shannon index, Chao1 index and species richness, as well as beta diversity, were estimated for the microbial communities using the vegan R package (25).

To predict the ecological interactions between each possible pair of the identified microbial species (*n* =ies, we created a half matrix of 124,251 species pair combinations (*n×*(*n*−1) *÷* 2). For each of those pairs, we applied flux balance analysis (FBA) using our self-developed R package EcoGS. This package employs metabolic modelling to compare the predicted growth of bacterial models with their co-growth after merging them into a community model using Sybil (26) and MicrobiomGS2 (27) R packages, respectively. The EcoGS package is available and described in more detail on GitHub (https://github.com/maringos/EcoGS) (28). To that end, we categorised each bacterial species pair into one of the possible six ecological interactions (supplementary table S2). Thereafter, we counted the number of pairs belonging to each ecological interaction in each microbial community and weighed those counts by the abundance of the bacteria in each pair. One effective way to account for the codependency within compositional data is to use a logarithmic transformation of the ratio between every two variables (each two ecological interaction percentages in our case). The six possible ecological interactions between bacteria resulted in 15 combinations of log ratios (log (Antagonism/Competition), log (Commensalism/Mutualism), … etc).

### 3.3. Microbial community modelling

We first filtered out the bacterial models that show low estimated growth (flux <10⁻³ on the modelled diet, flux units are arbitrary), as they can hinder community growth. Constraint-based modelling was implemented as flux balance analysis (FBA) using coupling constraints to simulate the community metabolic models using MicrobiomeGS (12) and in-house scripts. The bacterial models that belong to each single community were merged into a community model, where each bacterium has its own compartment and the community members exchange metabolites freely among each other and with the host in an environmental compartment. Thereafter, a community-level biomass reaction was introduced that accounts for the individual bacterial biomasses according to their relative abundance in the respective sample. The lower bounds of the exchange reactions were adjusted individually according to the estimated precision diet for each participant. An objective function was set to optimise for the community’s growth. The cplexAPI R package (29) was used to optimise for growth using the linear solver IBM ILOG CPLEX through its R interface with an academic license (30).

Thereafter, metabolites and reactions were filtered out if their fluxes had a low standard deviation (<0.01) among the different communities or if they were only present in less than 70% of the diabetic microbial communities.

### 3.4. Statistical analysis

The strong correlation between baseline glucose and HbA1c (P-value < 2.2e-16, rho = 0.96) led us to consider a particular measure in this study associated with blood glucose levels if it is correlated with either fasting glucose, HbA1c, or both.

All statistical analysis was conducted with R (31) within the Rstudio environment (32). The plots were created with the ggplot2 R package (33) and R base plotting functions. The results of all statistical tests were determined to be significant below a threshold of α=0.05. P-values were adjusted for multiple testing using false discovery rate correction (FDR). Confounders used for statistical test correction are: Gender, age, waist-to-hip ratio, activity, carbohydrate dietary intake and, if applicable, the antidiabetic medication treatment.

## 4. Results

### 4.1. Associations of individual species with diabetes with no differences in diversity measures

The comparison of bacterial abundance between diabetic and healthy individuals revealed three species from distinct genera (*Clostridium*, *Eubacterium* and *Lachnospiraceae*) exhibiting significantly higher abundances in healthy individuals. Conversely, diabetic patients showed higher abundances for eight species across five genera, including *Ruminococcus*, *Granulicatella*, *Clostridium*, *Bacteroides* and three species of *Streptococcus* as determined by the Mann-Whitney U test (Figure 1. A). Linear models that accounted for confounders (gender, age, waist-to-hip ratio, activity, and antidiabetic medication treatment) did not yield any significant association between blood glucose and alpha diversity indexes, neither for healthy individuals nor for people with diabetes. Although there is a significant increase in the alpha diversity measures in diabetic patients compared to healthy individuals, these differences disappear when correcting for the confounders, with the exception of the species richness, which was significantly increased in diabetic individuals. Moreover, no significant differences were observed between patients who are taking diabetes medications and those who are not using medication (Supplementary Table S3). Beta diversity, on the other hand, did not show any clear signal of differences between diabetic and healthy individuals.

**Figure 1:**
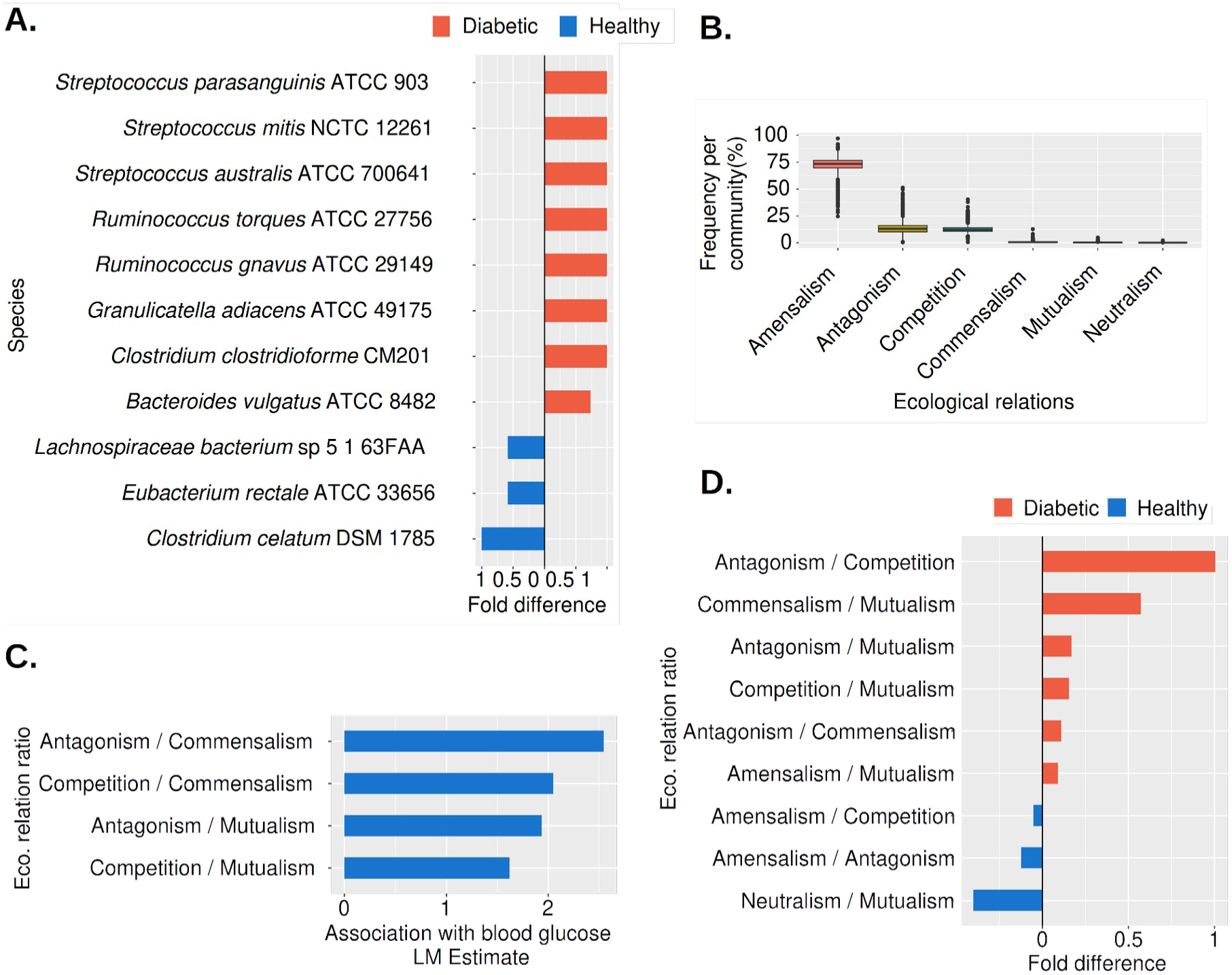
Microbial diversity and ecological interactions vs blood glucose and diabetes; *A.* Bacterial species with abundances that are significantly different between healthy and diabetic individuals, the fold difference was calculated as a log2 scale. *B.* The distribution of the frequencies for each of the six ecological interactions per microbial community. *C.* Ecological interaction ratios that are significantly associated with blood glucose. The estimate of the linear model is shown on the X-axis. *D.* Ecological interaction ratios that are significantly different between diabetic and healthy individuals. The fold difference of the ratios is shown on the X-axis in a log2 scale.

### 4.2. A relative increase in exploitative ecological interactions corresponding to glucose levels

Upon application of the EcoGS pipeline to the microbial communities, ecological relations were predicted for each pair of species within each community. These predictions were weighted by the abundance of the tested pair. Subsequently, the frequency of each type of ecological interaction within each community was estimated. Amensalistic ecological interactions accounted for the highest percentage of all ecological interactions, followed by antagonism competition, commensalism, mutualism and neutralism, respectively (Figure 1. B). Utilising linear models while controlling for confounders, a shift towards exploitative ecological interactions (antagonism and competition) at the expense of collaborative ecological interactions (mutualism and commensalism) was detected in association with increased glucose levels (Figure 1. C). Furthermore, a shift towards amensalism, antagonism, and competition (exploitative interactions) at the expense of mutualism (collaborative) was observed in diabetic individuals, along with an increase in antagonism (exploitative) relative to commensalism (collaborative). Meanwhile, in healthy individuals, amensalism was increased relative to competition and antagonism, as determined by the Mann-Whitney U test (Figure 1. D, Supplementary Figure S2).

### 4.3. Microbial metabolic fluxes are associated with blood glucose levels and diabetes

A total of 243 within-community fluxes and 243 metabolic fluxes between the community and the host were predicted using community flux balance analysis. The predicted fluxes with the host are either from the host’s diet to the community (diet-to-community fluxes) or from the community to the host (community-to-host fluxes). After filtering fluxes based on standard deviation and presence in diabetic samples, 105 within-community, 43 community-to-host, and 59 diet-to-community fluxes were tested for their association with host blood glucose levels. Linear regression analysis was used to assess the relationship between metabolic fluxes and blood glucose levels, considering confounding variables, namely sex, age, waist-to-hip ratio, physical activity and diabetic medication. Following multiple testing corrections (FDR), 71 metabolic fluxes were significantly different between diabetic patients and healthy individuals, using the Mann-Whitney U test with α=0.05 (Supplementary Figure S3). Those metabolites were grouped into 15 metabolic groups (Figure 2. A and Supplementary Table S4). Additionally, 31 metabolic fluxes were significantly associated with blood glucose levels (Figure 2. B).

**Figure 2:**
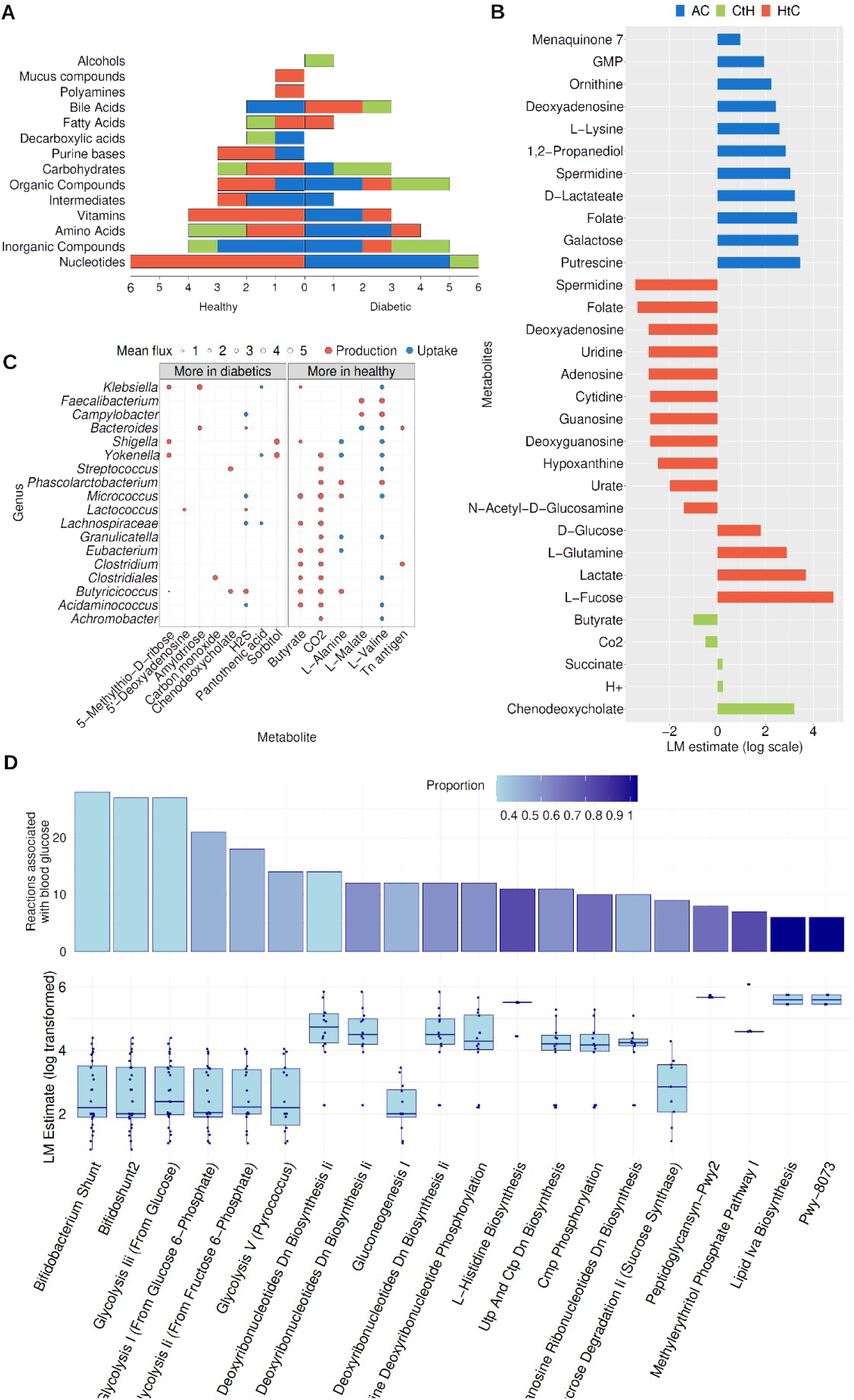
The simulated metabolic exchange fluxes vs. blood glucose levels and diabetes; *A.* The number of metabolites that are significantly different between diabetic and healthy individuals in each metabolic group, the actual metabolites are shown in supplementary figure S3. The type of metabolic exchange (within-community, community-to-host, and diet-to-community) are represented respectively in blue, red and green as stacked bars. Bars pointing to the right represent metabolites increased in healthy individuals, and those pointing to the left are increased in diabetic patients. *B.* The microbial metabolic fluxes that are significantly associated with the host blood glucose. Bars of negative associations point to the left, and bars of positive associations point to the right. The colours represent the three types of exchanges as in the previous plot A. The X-axis represents the estimated value of the linear model (coefficient of correlation). For visualisation purposes, those estimates were scaled as follows: log 10 ¿. *C.* Microbial production and uptake of metabolites that are associated with blood glucose levels. The average metabolic flux is on the X-axis, and the selected genera are on the Y-axis. The size of the circles represents the flux rate (log10 scale), with red dots for metabolites produced by the bacteria and blue dots for metabolites taken up by the bacteria. The left side shows metabolic fluxes that are higher in diabetic communities, and the right shows metabolic fluxes that are higher in healthy individuals. *D.* Enriched microbial metabolic subsystems. The upper bar plots represent the number of reactions associated with blood glucose in each subsystem, and the colour gradient of the bars reflects the proportion of the subsystem metabolites that are associated with blood glucose. The box plots in the lower part show the LM estimates for the reaction associations to blood glucose in each subsystem.

Investigating the fluxes of metabolites that are consumed by the community revealed that only four diet-to-community fluxes exhibited positive associations with blood glucose levels, namely D-glucose, glutamine, lactate and fructose. Conversely, the remaining eleven diet-to-community fluxes associated with blood glucose had negative associations. On the contrary, all of the eleven within-community fluxes associated with blood glucose had a positive association. Remarkably, among those metabolites, three have decreased diet-to-community fluxes while simultaneously increasing in their within-community fluxes accompanied with higher blood glucose. Those metabolites are: Spermidine, which has a role in alleviating oxidative stress and biofilm formation (34,35), folate which is necessary for nucleic acid biosynthesis, amino acid homeostasis, and methylation (36), (37) and 3-deoxyadenosine which is essential for energy metabolism, replication, and transcription.

Moreover, similarly to the association with blood glucose levels, multiple nucleotides, amino acids, carbohydrates, and vitamins were found to have significantly lower diet-to-community fluxes in diabetic patients. In contrast, nucleotides, amino acids, vitamins, and carbohydrates have increased within-community fluxes in diabetic patients (Figure 2. A and Supplementary Figure S3).

In summary, the metabolic fluxes among the microbiome are generally higher with increased blood glucose and in diabetic patients, and the metabolic fluxes from the diet to the microbiome are lower with increased blood glucose and in diabetic patients.

### 4.5. Dysbiosis in microbial metabolites provided to the host in diabetes

Three community-to-host metabolic fluxes were found to be positively associated with blood glucose, and two were found to be negatively associated (Fig 2. B). Notably, butyrate, a short-chain fatty acid, is produced at reduced levels by diabetic microbial communities (Fig. 2A). In addition, an increased production of H2S by the diabetic microbiome has been observed. Moreover, the community-to-host fluxes of L-valine and L-alanine are significantly lower in diabetic individuals. The same applies to the Tn antigen, which is a monosaccharide linked to serine or threonine and plays a role in mucin glycosylation (38).

Furthermore, the production and consumption of metabolites associated with diabetes by individual genera was investigated. This was done for 18 genera, comprising the five most abundant genera in the communities, along with the genera that showed a significant difference between healthy and diabetic individuals. (Figure 2. C). Among the tested genera, there are nine butyrate producers, including *Clostridium*, *Lachnospiraceae* and *Eubacterium.* Those Butyrate producers were found to have higher abundances in healthy individuals. Other genera include *Faecalibacterium* and *Campylobacter,* which are found to produce malate and valine, and three genera that are found to produce Alanine, which are *Phascolarctobacterium*, *Micrococcus,* and *Butyricicoccus*. And two genera produce the Tn antigen.

### 4.6. Nucleotide, bile acid, B vitamins and amino acids exhibit altered fluxes in the diabetic microbiome

Six out of the eleven diet-to-community metabolites that are negatively associated with blood glucose are nucleotides, namely, deoxyadenosine, adenosine, uridine, cytidine, deoxyguanosine, and guanosine. The flux for those six metabolites from the host to the community is also significantly decreased in diabetic patients. In contrast, the fluxes of guanosine monophosphate and deoxyadenosine within the community are positively associated with blood glucose. And in diabetic patients, the fluxes of deoxyadenosine, cytidine, uridine, guanosine monophosphate, and uridine monophosphate are increased within the community. Additionally, in diabetic patients, there is an elevated flux of deoxyadenosine from the community to the host. Nucleotide metabolism plays crucial roles in energy metabolism, nucleic acid synthesis and repair, cell-to-cell communication and signal transduction in bacteria (39). Nucleotides are also utilised in most metabolic pathways for energy exchange and regulation (40). It is noteworthy that bile acids follow an opposite trend, where the diet-to-community fluxes of taurochenodeoxycholate and taurocholate increase in diabetes. And those bile acids are deconjugated by the microbiome to cholate and taurine. The community-to-host fluxes of chenodeoxycholate are also increased in diabetes. Meanwhile, the within-community fluxes of cholate and 3-dehydrocholate are decreased in diabetes.

Furthermore, among diabetic individuals, B-vitamins such as nicotinamide (B3), riboflavin (B2), thiamin (B1), and folate (B9) exhibit decreased diet-to-community fluxes, excluding Niacin (another form of B3), which increases in diabetes. Meanwhile, vitamin K, in two forms, also increased within-community flux in diabetic microbiomes.

Similarly, the diet-to-community fluxes of amino acids, L-tryptophan and L-tyrosine are reduced in diabetic patients, while the flux of L−glutamine is increased. The amino acids taurine (potentially derived from the conjugated bile acids), ornithine, and L-valine have increased within-community fluxes, while L−valine and L−alanine have increased community-to-host fluxes in diabetic individuals.

### 4.7. Energy production and core metabolic pathways are positively associated with elevated blood glucose

We summarised all the internal reactions from all bacteria within each microbial community and tested their association with the blood glucose levels of the host. Starting with 4,283 reactions, after filtration as described in the methods, we ended up with 855 reactions with a maximum of 840, a minimum of 517, and a median of 796 internal reactions per microbial community. By applying a linear model with confounders as described in the methods, we detected 507 reactions that are significantly associated with blood glucose; each of these reactions belongs to one or multiple of 415 pathways (subsystems). Thereafter, we carried out an enrichment analysis using Fisher’s exact test, resulting in 20 pathways that are significantly enriched with reactions that are associated with blood glucose. Most of those pathways belong to core metabolism, with the majority involved in energy metabolism.

We illustrated six major pathways in (Figure 3) based on EcoCyc metabolic maps (41) to observe connections among the enriched pathways and with the detected metabolites. Among the enriched pathways;

● **Glycolysis Ⅰ**, which presents the first step in energy production from D-glucose by splitting it into pyruvates. This pathway results in the formation of ATP along with twelve intermediates that can be used as carbon skeletons for biosynthesis. Pyruvate in turn is used in **amino acid Ⅰ synthesis**, **gluconeogenesis** as well in **mixed acid fermentation.**
● **Gluconeogenesis I** results in the biosynthesis of glucose from other non-carbohydrate sources, playing a role in energy production and regulation, and its balance with **glycolysis** maintains energy homeostasis in bacteria.
● **Mixed acid fermentation** also has an important role in energy production. Glucose or other sugars are broken down anaerobically to produce various acids while generating ATP and other compounds such as a-menaquinone, succinate, and malate. These can, in turn, be used by **Gluconeogenesis I** to produce D-glucose and maintain homeostasis.
● **Calvin-Benson-Bassham cycle** is a central pathway that is also involved in energy production by using high-energy molecules to drive the production of glyceraldehyde-3-phosphate as well as β-D-fructofuranose-6-phosphate that enters the **Starch biosynthesis** pathway to produce starch.
● **Sucrose degradation IV** (sucrose phosphorylase), another pathway involved in energy metabolism, also produces β-D-fructofuranose 6-phosphate, which can enter the **Bifidobacterium shunt.**
● **Bifidobacterium shunt** in turn produces lactate, which can be used by butyrate-producing bacteria like some of the *Clostridium* genus to produce acetate and butyrate(42). Moreover, this pathway can produce D-ribose 5-phosphate which is essential for nucleotide biosynthesis.

**Figure 3:**
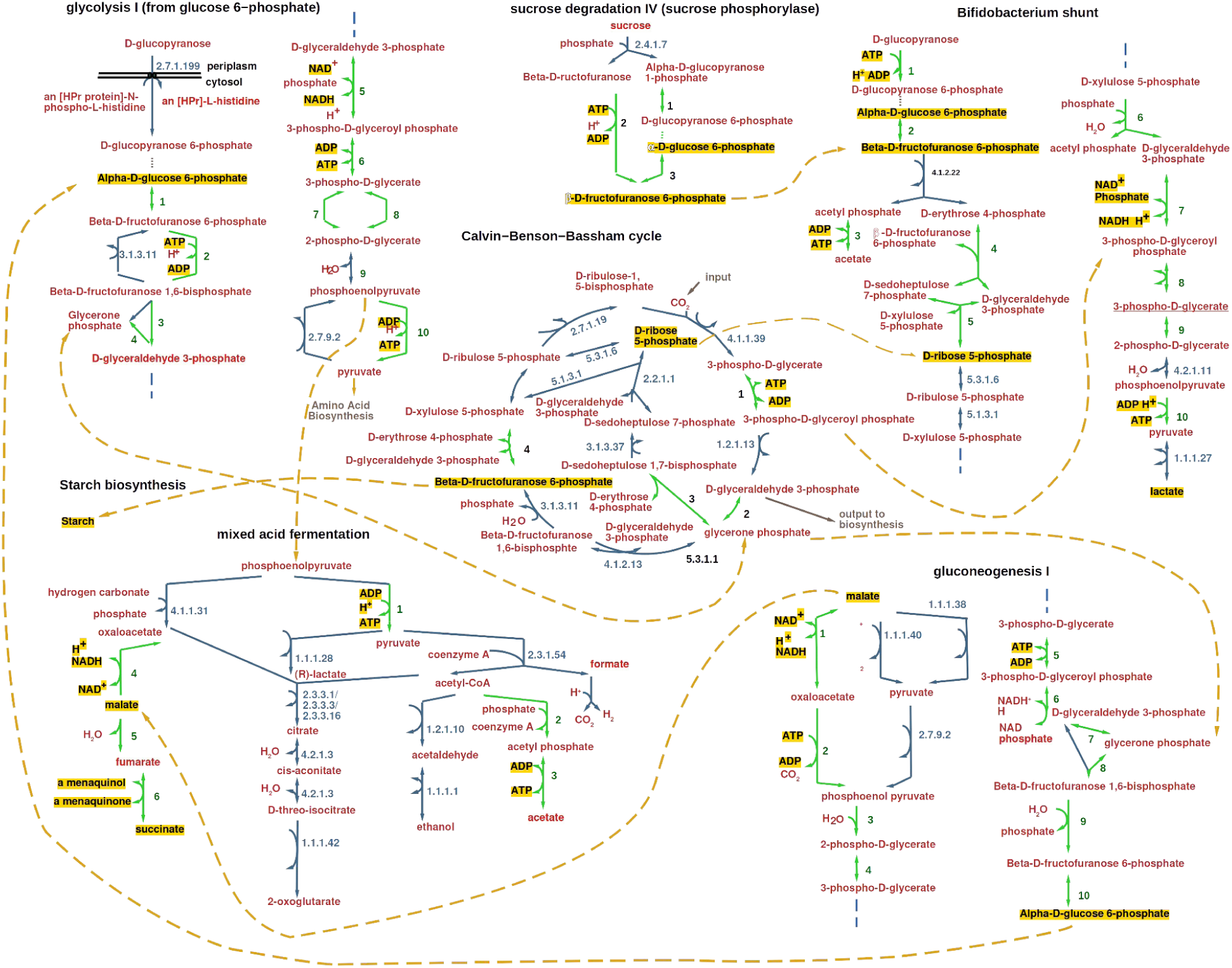
Metabolic maps of subsystems enriched with reactions associated with blood glucose. An example of six major core metabolism pathways enriched with microbial reactions positively associated with the host’s blood glucose. Arrows represent reactions, and two-sided arrows are used for reversible reactions. Green arrows represent reactions that are significantly associated with blood glucose, while blue arrows represent reactions that are not. Numbers next to green arrows are described in supplementary table S5, while the ID numbers next to blue arrows are KEGG identifiers (43). The metabolites are in red, and metabolites with fluxes that are associated with blood glucose, as well as those known to have connections to diabetes, are in black text with yellow highlights. Dashed orange arrows connect the pathways through shared metabolites.

## 5. Discussion

In this study, we conducted a metabolic modelling-based analysis of changes in the gut microbiome associated with increased blood glucose levels and diabetes. This included alterations in the interactions within the gut microbial community and between the community and the human host. To this end, we acquired 16S sequences along with metadata and dietary information from 1,866 individuals, including 81 diabetic patients.

It is important to highlight that participants in this study were not selected randomly; rather, they were individuals with a general interest in enhancing their health through personalised dietary recommendations (such as to lose weight or improve general fitness), which is a core aspect of Perfood’s service. We corrected for confounders in the statistical tests as it is known that the gut microbiome can be influenced by multiple factors, including diet, lifestyle, and medication (44). Moreover, by accounting for diabetes medication and carbohydrate intake in the regression analysis, we aimed to ensure that the associations between the microbiome metabolic functions and the blood glucose were independent of the influence of other environmental parameters.

The differential abundance analysis results are consistent with previous reports of an increase in *Streptococcus* and *Bacteroides* and a decrease in *Clostridium* among other butyrate-producing genera in diabetic children (45) and women (46). Multiple studies have reported that diabetes is usually accompanied by reduced gut microbial diversity (47,48). In this study, the observed increased richness in diabetic individuals with no difference in the other alpha diversity measures. This indicates that diabetic communities have a higher number of species but with low abundance, not resulting in an increased alpha diversity (Table S3). However, the simple measures of microbial diversity are limited in representing the actual changes in the community dynamics. Taxonomic diversity measures are unaware of the diversity on the genetic and metabolic levels; thus, they are susceptible to varying observations across cohorts (49,50).

The ecological interaction predictions in this study revealed a relative decrease in collaborative interactions (mutualism and commensalism) and an increase in exploitative interactions (antagonism and competition) in diabetic microbial communities. Such dysbiosis is potentially promoted by the high availability of simple energy sources such as glucose. This observation is consistent with the associations observed in a separate study, where an individual’s dietary intake of glucose is linked to a decrease in mutualism and an increase in antagonism and competition within their gut microbiome.

Our investigation revealed extensive associations between blood glucose levels and the microbiome metabolic functions as well as between clinically diagnosed diabetes and the microbiome metabolic functions. Notably, all the metabolic fluxes that are positively associated with blood glucose were found to be significantly higher in diabetic patients, while negatively associated metabolites were significantly lower in diabetic patients, which provides a proof of concept. Elevated blood glucose was accompanied by increased fluxes of eleven metabolites among the microbial community members. Also, eleven metabolites showed a decrease in their flux from the host’s diet to the microbial community (Figure 2. B). Among those metabolites, nucleotides, B vitamins, and amino acids had elevated fluxes within the community and decreased fluxes from the host to the community with higher blood glucose levels. This supports the notion of a diabetic microbiome that uses the vastly available glucose from the environment and becomes less controlled by the host and more dependent on the metabolic interchanges among its members.

With higher blood glucose, four metabolites, D-glucose, glutamine, lactate and fructose, showed higher diet-to-community flux. Those metabolites provide energy for the community and substantiate the connection between blood glucose levels and glucose availability to the gut microbiome. This agrees with the previously reported increase in the amounts of faecal carbohydrates, especially monosaccharides, in individuals displaying insulin resistance, where the elevated levels of monosaccharides were found to have a direct link to microbial carbohydrate metabolisms (18).

Moreover, the observed reduction in the production of the short-chain fatty acid butyrate in the diabetic community is consistent with previous studies reporting a decreased abundance of butyrate-producing bacteria in diabetic patients (45,47,48,51). In turn, it coincides with reported decreased faecal butyrate concentrations in type 1 diabetic patients in comparison with healthy controls (52). Butyrate is also known to have an important role in moderating inflammatory responses (53,54). In addition, it constitutes an essential source of energy for the colon and plays a signalling role by binding to G-protein coupled receptor (Gpr41 and Gpr43), which suppresses insulin signalling and prevents the accumulation of fat; therefore, butyrate helps to maintain the energetic balance (55). A reduction in the diet-to-community flux of L-tryptophan was also observed in this study. Tryptophan can be converted by the bacteria into indole that acts as a signalling molecule and plays a role in regulating the integrity of the intestinal barrier, hence influencing liver health and systemic metabolic balance (56), in addition to its anti-inflammatory role mediated by AhR signalling. Moreover, both tryptophan and indole have been reported to have an important impact on inflammation and insulin sensitivity (57,58). In addition, the increased production of H2S by the diabetic microbiome aligns with a report by Quin and others of an increase in sulfate-reducing bacteria in the diabetic microbiome (51). It is important to note that in normal amounts, H2S plays essential roles in gut motility, sensing, secretion, and absorption, suggesting potential implications for gastrointestinal disease therapies. Another metabolite that was increased in diabetic microbial communities is sorbitol, which is produced by *Yokenella* and *Shigella* and was previously reported to have glucose intolerance-inducing effects (59). All that make butyrate, Tryptophan and H2S potential targets for intervention-based treatment of diabetes.

B vitamins that exhibit decreased diet-to-community fluxes in diabetic individuals act as coenzymes in various cellular reactions. They participate in all energy-producing processes within cells (60,61). Despite the reduced uptake of these vitamins by the community, it is noteworthy that multiple bacteria can synthesise B vitamins. This, coupled with the heightened energy metabolism in diabetic gut communities, supports the notion of a lesser degree of control by the host’s diet over the diabetic microbiomes.

Notably, all of the associations between the microbial internal reactions and blood glucose were positive. This is consistent with the observation that all the metabolic exchange fluxes among the community (within-community fluxes) are also positively associated with blood glucose (Figure 2.D). The observed enriched microbial pathways in diabetes are essential for energy metabolism, their increased activity in correspondence to high blood glucose levels further supports the notion that the microbiome becomes active in producing the energy it needs for its growth, starting with glycolysis that utilises glucose to the gluconeogenesis that maintains homeostasis.

## 6. Concluding remarks

In light of our observations, we concluded that the availability of glucose as a simple source of energy, along with the disturbance of the community structure caused by diabetes, alters the metabolic relationship between the host and microbiome, causing a state of dysbiosis. The diabetic microbiome becomes less dependent on the host, less shaped by the host’s diet, more active in its energy production, and more hostile and has more exploitative interactions among its member species.

Our study emphasises the necessity for comprehensively exploring the intricate interplay between diabetes and the gut microbial community. It provides an important basis for further experimental validations and dietary intervention analysis to explore how to potentially alter the dysbiotic effects of diabetes on the interactions between the microbiome and the host.

## 7. Ethics statement

The 16S rRNA sequencing data was generated in the context of a prior project (University of Lübeck, ethics notification ID 20-415). All patients gave their consent to the general use of their anonymised data for scientific purposes. Sequencing data and metadata were provided completely anonymised and thus did not require separate ethical approval according to national regulations for the analysis presented in this manuscript.

## 8. Acknowledgement

We acknowledge support by the German Research Foundation within the framework of the Excellence Cluster “Precision Medicine in Chronic Inflammation” (project code EXC2167).

We thank Jan Taubenheim, Karlis Arturs Moors, Silvio Washina, Johannes Zimmermann, and Robin Koch for fruitful discussion and critical comments.

## 9. Author Contributions

Conceptualisation, ASK, CK, AK, TS; Methodology, ASK, CK; Software, ASK, GM; Data Collection and curation, AK, TS, JZ, OW; Data Analysis, ASK; Writing the manuscript, ASK with revision by all coauthors, Visualization, ASK; Funding Acquisition, CK, TS; Project Administration, CK.

## 10. Conflict of interests

The co-authors, Axel Künstner, Torsten Schröder, Julius Zauleck, and Oliver Witt, are affiliated with Perfood GmbH. All authors declare no competing or commercial interests.

The data for this study were collected up to 2021, and the microbiome data is not part of any product by Perfood GmbH as of today.

## 11. Supplementary

**Figure S1:**
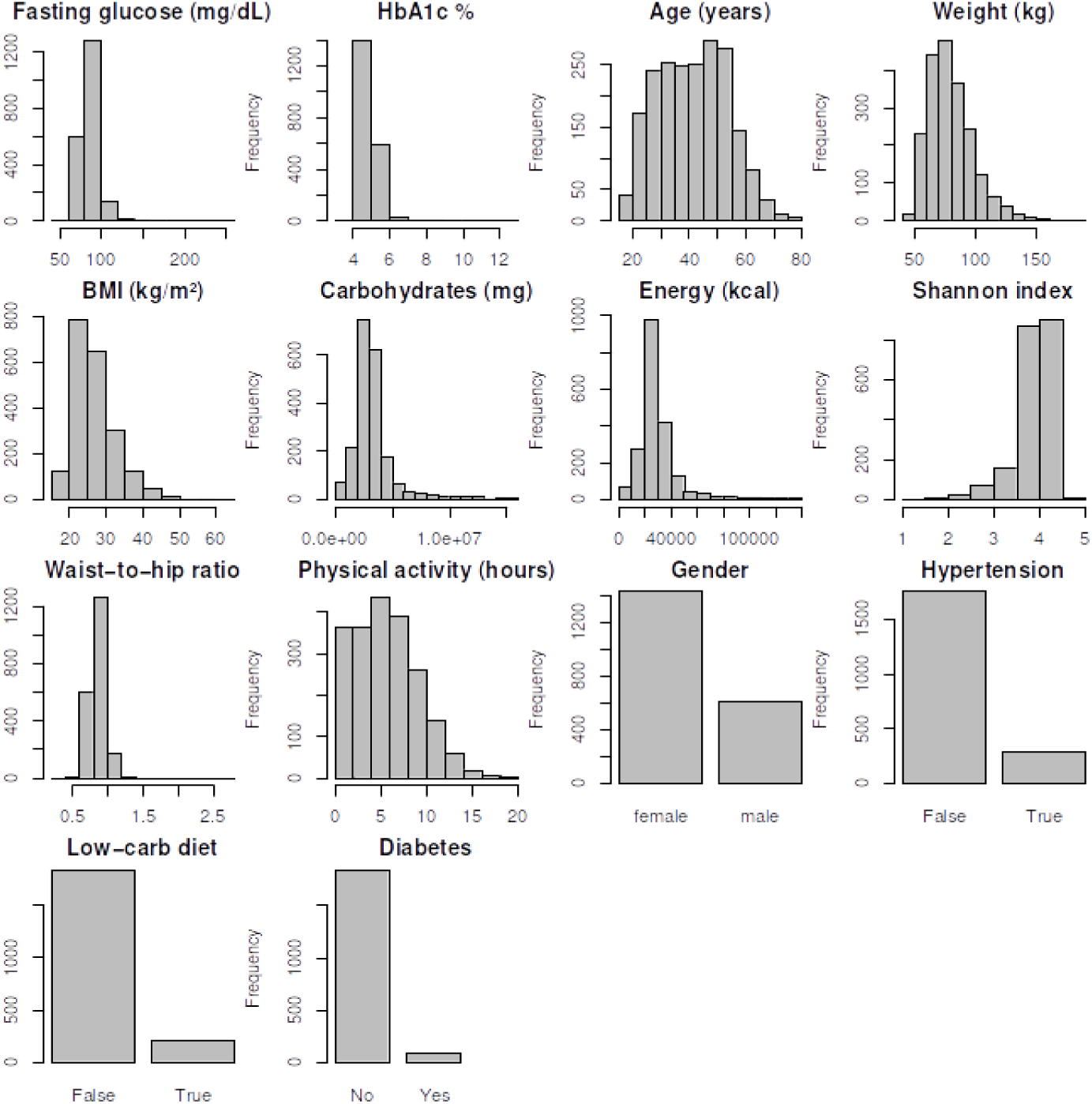
Histogram distributions explaining the metadata.

**Table S1:** dada2 package functions used for the 16s sequence preparation.

|  |  |
| --- | --- |
| learnErrors | Used for estimating error rates (kmer based). |
| plotErrors | Provides a graphical representation of the error rate. |
| filterAndTrim | Refines sequencing data by filtering out low-quality reads and trimming sequences. |
| derepFastq | Removes duplicate sequences. |
| dada | Used for denoising sequencing data and resolving unique variants. |
| mergePairs | Aligns and merges paired-end sequencing reads. |

**Table S2:** Ecological interaction types and categories.

| Species A | Species B | Ecological interaction | Interaction Category |
| --- | --- | --- | --- |
| Benefit | Benefit | Mutualism | Collaborative |
| No change | Benefit | Commensalism | Collaborative |
| Benefit | Harm | Antagonism | Exploitative |
| No change | No change | Neutralism | Neutral |
| No change | Harm | Amensalism | Exploitative |
| Harm | Harm | Competition | Exploitative |

**Table S3:** Alpha diversity measures vs. diabetes, fasting glucose, and medications anti-diabetes. * significantly increased alpha diversity with the observation. LM: Linear models with confounders All P-values are adjusted for false discovery rates (FDR correction)

|  | Diabetes |  | Higher<br>fasting<br>glucose | Anti-diabetic medications |  |
| --- | --- | --- | --- | --- | --- |
|  | Mann-Whitney, FDR P-values | LM, FDR P-values | LM, FDR P-values | Mann-Whitney, FDR P-values | LM, FDR P-values |
| Shannon | $5.4 \times 10^{-6}$ * | 0.051 | 0.08 | 0.44 | 0.37 |
| Chao1 | $2.1 \times 10^{-4}$ * | 0.14 | 0.08 | 0.45 | 0.62 |
| Species richness | $3.7 \times 10^{-5}$ * | 0.003 * | 0.32 | 0.49 | 0.37 |
| Simpson's | $1.1 \times 10^{-3}$ * | 0.5 | 0.09 | 0.45 | 0.37 |

**Figure S2:**
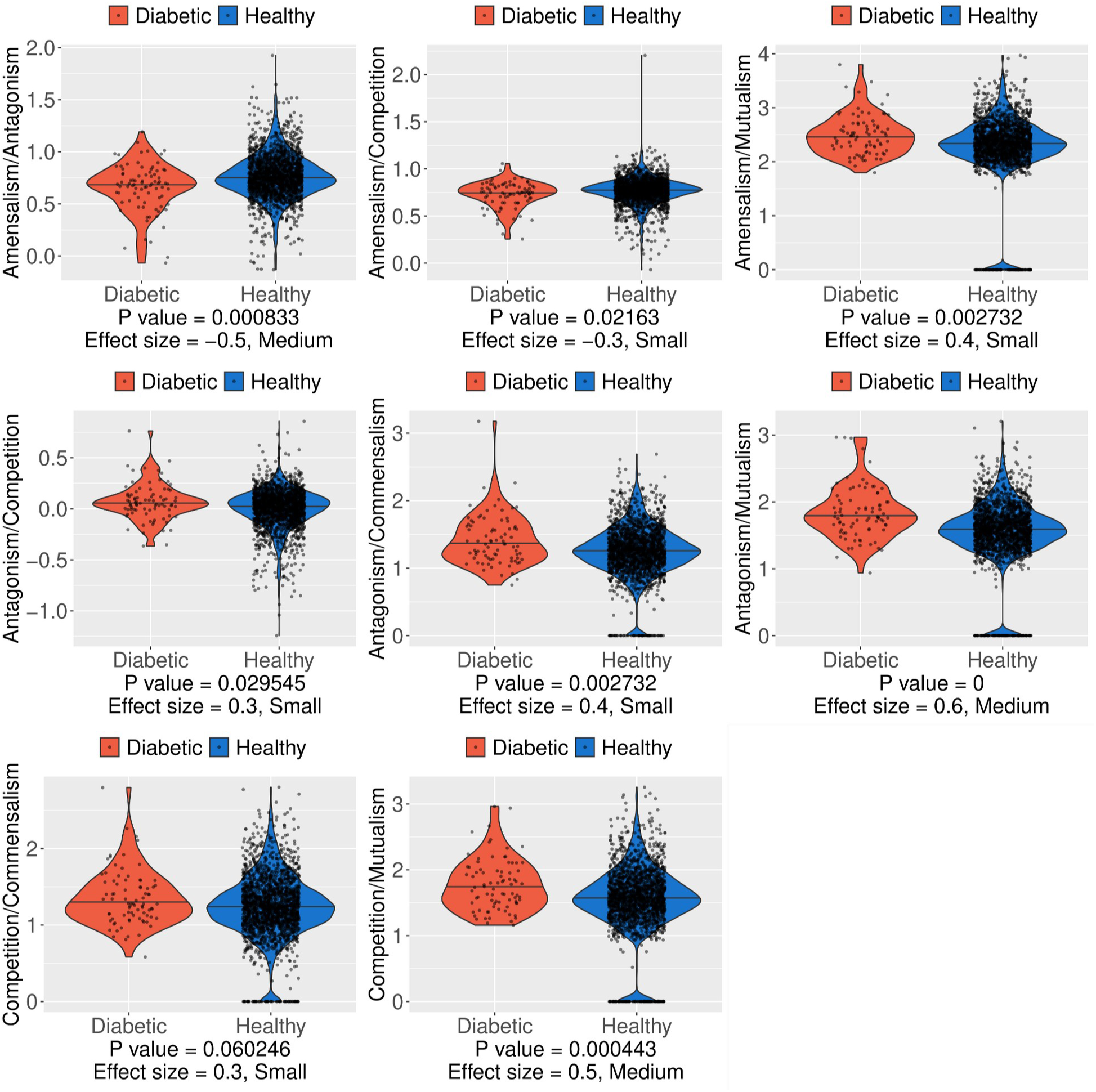
The ecological interaction rations between diabetic and non-diabetic microbial communities, Mann-Whitney U test.

**Figure S3:**
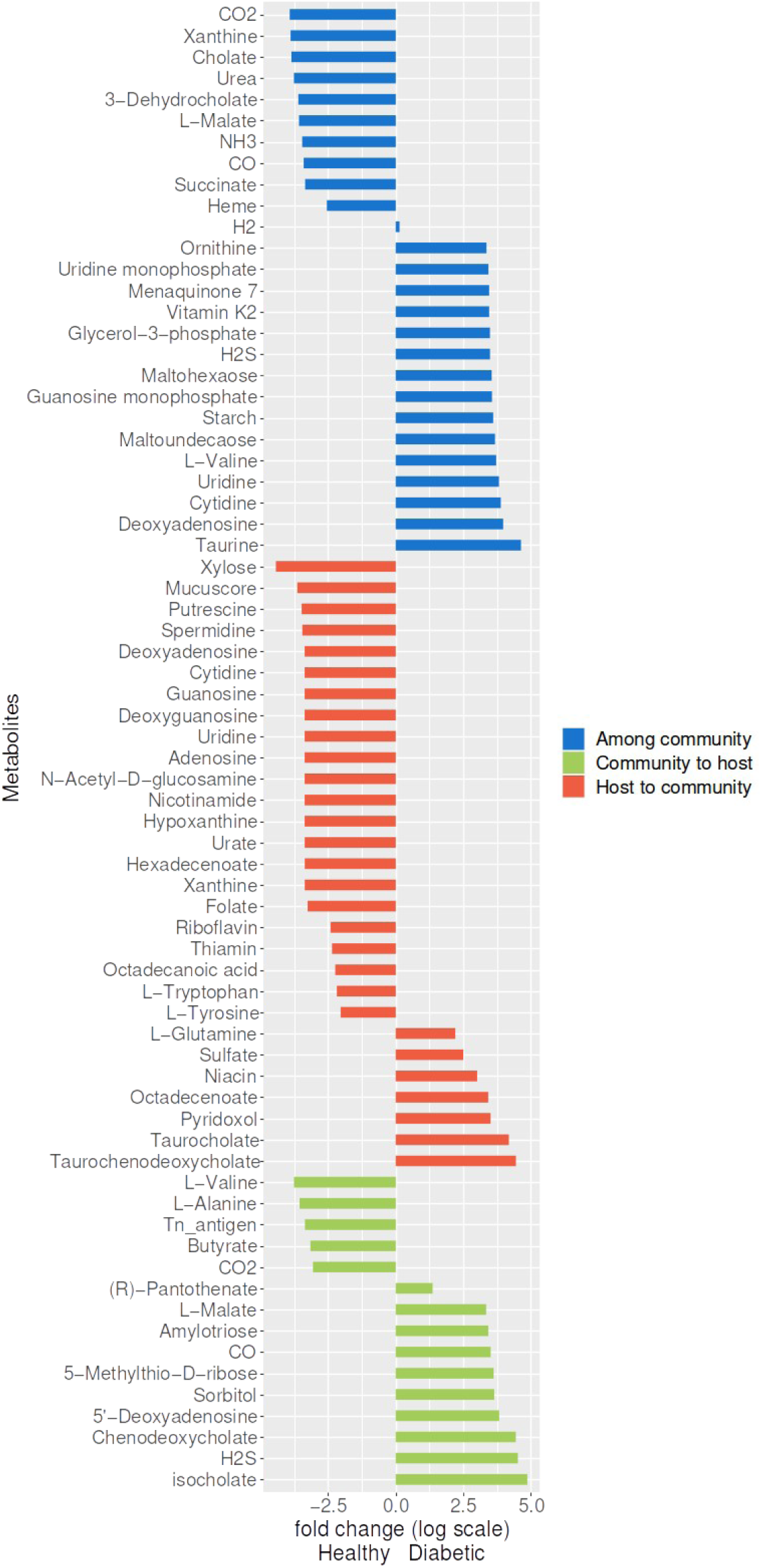
Metabolic fluxes associated with diabetes, to the left increased in healthy and to the right increased in diabetic.

**Table S5:** The pathways and their reactions that are positively associated with blood glucose.

|  |
| --- |
| **Glycolysis I (from glucose 6–phosphate), ten reactions:** 1) glucose-6-phosphate isomerase, 2) 6-phosphofructokinase, 3) fructose-bisphosphate aldolase, 4) triose-phosphate isomerase, 5) glyceraldehyde-3-phosphate dehydrogenase, 6) phosphoglycerate kinase, 7) 2,3-bisphosphoglycerate-independent phosphoglycerate mutase, 8) phosphoglycerate mutase, 9) enolase, 10) pyruvate kinase. |
| **Gluconeogenesis I, nine reactions;** 1) malate dehydrogenase, 2) 2- phosphoenolpyruvate carboxykinase, 3) enolase, 4) 2,3-bisphosphoglycerate-independent phosphoglycerate mutase, 5) phosphoglycerate kinase, 6) glyceraldehyde-3-phosphate dehydrogenase, 7) triose-phosphate isomerase, 8) fructose-bisphosphate aldolase, 9) Missing reaction: fructose-1,6-bisphosphatase, 10) glucose-6-phosphate isomerase |
| **Calvin–Benson–Bassham cycle, four reactions:** 1)Phosphoglycerate kinase, 2)triosephosphate isomerase, 3)sedoheptulose-1,7-bisphosphate aldolase, 4)D-fructose, 6)phosphate: D-glyceraldehyde-3-phosphate glycolaldehydetransferase |
| **Bifidobacterium shunt, ten reactions;** 1)ATP: D-glucose 6-phosphotransferase, 2)Glucose-6 phosphate isomerase, 3)ATP: acetate phosphotransferase, 4)sedoheptulose-7-phosphate, 5)D-Fructose 6-phosphate, 6)xylulose-5-phosphate phosphoketolase, 7)D-glyceraldehyde-3-phosphate, 8)ATP:3-phospho-D-glycerate 1-phosphotransferase, 9)2-Phospho-D-glycerate 2,3-phosphomutase, 10)ATP: pyruvate 2-O-phosphotransferase. |
| **Mixed acid fermentation, six reactions:** 1) pyruvate kinase, 2) phosphate acetyltransferase, 3) acetate kinase, 4) malate dehydrogenase, 5) fumarase, 6) fumarate reductase |
| **Sucrose degradation IV (sucrose phosphorylase), three reactions:** 1) phosphoglucomutase, 2) fructokinase, 3) glucose-6 phosphate isomerase |

## References

1. Xie J, Wang M, Long Z, Ning H, Li J, Cao Y, et al. Global burden of type 2 diabetes in adolescents and young adults, 1990-2019: systematic analysis of the Global Burden of Disease Study 2019. Br Med J. 2022 Dec 7;379:e072385.

2. Rinninella E, Raoul P, Cintoni M, Franceschi F, Miggiano GAD, Gasbarrini A, et al. What is the Healthy Gut Microbiota Composition? A Changing Ecosystem across Age, Environment, Diet, and Diseases. Microorganisms. 2019 Jan 10;7(1):14.

3. Cunningham AL, Stephens JW, Harris DA. Gut microbiota influence in type 2 diabetes mellitus (T2DM). Gut Pathog. 2021 Aug 6;13(1):50.

4. Pan Y, Bu T, Deng X, Jia J, Yuan G. Gut microbiota and type 2 diabetes mellitus: a focus on the gut-brain axis. Endocrine. 2021 Jan 16;84:1–15.

5. Ghosh SS, Wang J, Yannie PJ, Ghosh S. Intestinal Barrier Dysfunction, LPS Translocation, and Disease Development. J Endocr Soc. 2020 Feb 1;4(2):bvz039.

6. Herrema H, Niess JH. Intestinal microbial metabolites in human metabolism and type 2 diabetes. Diabetologia. 2020 Dec 1;63(12):2533–47.

7. Gurung M, Li Z, You H, Rodrigues R, Jump DB, Morgun A, et al. Role of gut microbiota in type 2 diabetes pathophysiology. eBioMedicine. 2020 Jan 1;51(2020):102590.

8. Michellod D, Liebeke M. Host–microbe metabolic dialogue. Nat Microbiol. 2024 Feb 5;9(2):318–9.

9. Zimmermann J, Kaleta C, Waschina S. gapseq: informed prediction of bacterial metabolic pathways and reconstruction of accurate metabolic models. Genome Biol. 2021 Mar 10;22(1):81.

10. Zacharias HU, Kaleta C, Cossais F, Schaeffer E, Berndt H, Best L, et al. Microbiome and Metabolome Insights into the Role of the Gastrointestinal–Brain Axis in Parkinson’s and Alzheimer’s Disease: Unveiling Potential Therapeutic Targets. Metabolites. 2022 Dec;12(12):1222.

11. Forero-Rodríguez LJ, Josephs-Spaulding J, Flor S, Pinzón A, Kaleta C. Parkinson’s Disease and the Metal–Microbiome–Gut–Brain Axis: A Systems Toxicology Approach. Antioxidants. 2022 Jan;11(1):71.

12. Aden K, Rehman A, Waschina S, Pan WH, Walker A, Lucio M, et al. Metabolic Functions of Gut Microbes Associate With Efficacy of Tumor Necrosis Factor Antagonists in Patients With Inflammatory Bowel Diseases. Gastroenterology. 2019 Nov 1;157(5):1279–1292.e11.

13. Aden K, Rehman A, Pan WH, Waschina S, Barthi R, Zimmermann J, et al. The gut microbiome in IBD is characterized by impaired metabolic cooperativity and can be restored upon anti-TNFα therapy. In: Zeitschrift für Gastroenterologie. Georg Thieme Verlag KG; 2017. p. KV 135.

14. Ezzamouri B, Rosario D, Bidkhori G, Lee S, Uhlen M, Shoaie S. Metabolic modelling of the human gut microbiome in type 2 diabetes patients in response to metformin treatment. NPJ Syst Biol Appl. 2023 Jan 21;9.

15. Väremo L, Nookaew I, Nielsen J. Novel insights into obesity and diabetes through genome-scale metabolic modeling. Front Physiol. 2013 Apr 25;4.

16. Proffitt C, Bidkhori G, Lee S, Tebani A, Mardinoglu A, Uhlen M, et al. Genome-scale metabolic modelling of the human gut microbiome reveals changes in the glyoxylate and dicarboxylate metabolism in metabolic disorders. iScience. 2022 Jun 2;25(7):104513.

17. Jeckelmann JM, Erni B. Transporters of glucose and other carbohydrates in bacteria. Pflüg Arch – Eur J Physiol. 2020 Sep 1;472(9):1129–53.

18. Takeuchi T, Kubota T, Nakanishi Y, Tsugawa H, Suda W, Kwon ATJ, et al. Gut microbial carbohydrate metabolism contributes to insulin resistance. Nature. 2023 Sep;621(7978):389–95.

19. Twesten C, Schröder T, Leitow FM. Improved method for determining blood glucose responses [Internet]. EP3896700A1, 2021. Available from: https://patents.google.com/patent/EP3896700A1/de?assignee=perfood

20. Kordowski A, Tetzlaff-Lelleck VV, Speckmann B, Loh G, Künstner A, Schulz F, et al. A nutritional supplement based on a synbiotic combination of Bacillus subtilis DSM 32315 and L-alanyl-L-glutamine improves glucose metabolism in healthy prediabetic subjects – A real-life post-marketing study. Front Nutr. 2022 Dec 8;9.

21. Callahan BJ, McMurdie PJ, Rosen MJ, Han AW, Johnson AJA, Holmes SP. DADA2: High-resolution sample inference from Illumina amplicon data. Nat Methods. 2016 Jul;13(7):581– 3.

22. Pryor R, Norvaisas P, Marinos G, Best L, Thingholm LB, Quintaneiro LM, et al. Host-Microbe-Drug-Nutrient Screen Identifies Bacterial Effectors of Metformin Therapy. Cell. 2019 Sep 5;178(6):1299–1312.e29.

23. Magnúsdóttir S, Heinken A, Kutt L, Ravcheev DA, Bauer E, Noronha A, et al. Generation of genome-scale metabolic reconstructions for 773 members of the human gut microbiota. Nat Biotechnol. 2017 Jan;35(1):81–9.

24. Edgar RC. Search and clustering orders of magnitude faster than BLAST. Bioinformatics. 2010 Oct 1;26(19):2460–1.

25. Oksanen J, Simpson G, Blanchet F, Kindt R, Legendre P, Minchin P, et al. vegan: Community Ecology Package_. R package version 2.6-4. [Internet]. 2022. Available from: <https://CRAN.R-project.org/package=vegan>

26. Gelius-Dietrich G, Desouki AA, Fritzemeier CJ, Lercher MJ. sybil – Efficient constraint-based modelling in R. BMC Syst Biol. 2013 Nov 13;7(1):125.

27. Waschina S. Analysis and simulation of genome-& ecosystem-scale microbial metabolism [Internet]. Available from: www.github.com/Waschina/MicrobiomeGS2.

28. Kadibalban S, Marinos G, Moors K, Waschina S, Kaleta C. EcoGS, Available from: https://github.com/maringos/EcoGS.

29. Gelius-Dietrich G. cplexAPI: R Interface to C API of IBM ILOG CPLEX [Internet]. Available from: https://CRAN.R-project.org/package=cplexAPI.

30. Cplex, I.I. (2009). V12.1: User’s Manual for CPLEX. International Business Machines Corporation, 46(53), p.157.

31. R Core Team. R: A Language and Environment for Statistical Computing [Internet]. Vienna, Austria: R Foundation for Statistical Computing; Available from: https://www.R-project.org/.

32. Posit team (2023). RStudio: Integrated Development Environment for R. Posit Software, PBC, Boston, MA. URL http://www.posit.co/.

33. Wickham H (2016). ggplot2: Elegant Graphics for Data Analysis. Springer-Verlag New York. ISBN 978-3-319-24277-4, https://ggplot2.tidyverse.org.

34. Thongbhubate K, Nakafuji Y, Matsuoka R, Kakegawa S, Suzuki H. Effect of Spermidine on Biofilm Formation in Escherichia coli K-12. J Bacteriol. 2021 Apr 21;203(10):10.1128/jb.00652-20.

35. Kumar V, Mishra RK, Ghose D, Kalita A, Dhiman P, Prakash A, et al. Free spermidine evokes superoxide radicals that manifest toxicity. Wade JT, Storz G, Wade JT, editors. eLife. 2022 Apr 13;11:e77704.

36. Mölzer C, Wilson HM, Kuffova L, Forrester JV. A Role for Folate in Microbiome-Linked Control of Autoimmunity. J Immunol Res. 2021 May 20;2021:e9998200.

37. Kok DE, Steegenga WT, Smid EJ, Zoetendal EG, Ulrich CM, Kampman E. Bacterial folate biosynthesis and colorectal cancer risk: more than just a gut feeling. Crit Rev Food Sci Nutr. 2020 Jan 19;60(2):244–56.

38. Coletto E, Savva GM, Latousakis D, Pontifex M, Crost EH, Vaux L, et al. Role of mucin glycosylation in the gut microbiota-brain axis of core 3 O-glycan deficient mice. Sci Rep. 2023 Aug 26;13:13982.

39. Zakataeva NP. Microbial 5′-nucleotidases: their characteristics, roles in cellular metabolism, and possible practical applications. Appl Microbiol Biotechnol. 2021 Oct 1;105(20):7661–81.

40. Kilstrup M, Hammer K, Ruhdal Jensen P, Martinussen J. Nucleotide metabolism and its control in lactic acid bacteria. FEMS Microbiol Rev. 2005 Aug 1;29(3):555–90.

41. Keseler IM, Gama-Castro S, Mackie A, Billington R, Bonavides-Martínez C, Caspi R, et al. The EcoCyc Database in 2021. Front Microbiol. 2021 Jul 28;12:711077.

42. Detman A, Mielecki D, Chojnacka A, Salamon A, Błaszczyk MK, Sikora A. Cell factories converting lactate and acetate to butyrate: Clostridium butyricum and microbial communities from dark fermentation bioreactors. Microb Cell Factories. 2019 Feb 13;18(1):36.

43. Kanehisa M, Goto S. KEGG: kyoto encyclopedia of genes and genomes. Nucleic Acids Res. 2000 Jan 1;28(1):27–30.

44. Zhang S, Cai Y, Meng C, Ding X, Huang J, Luo X, et al. The role of the microbiome in diabetes mellitus. Diabetes Res Clin Pract. 2020 Dec 24;172:108645.

45. de Goffau MC, Fuentes S, van den Bogert B, Honkanen H, de Vos WM, Welling GW, et al. Aberrant gut microbiota composition at the onset of type 1 diabetes in young children. Diabetologia. 2014 Aug 1;57(8):1569–77.

46. Karlsson FH, Tremaroli V, Nookaew I, Bergström G, Behre CJ, Fagerberg B, et al. Gut metagenome in European women with normal, impaired and diabetic glucose control. Nature. 2013 Jun;498(7452):99–103.

47. Hu YH, Meyer K, Lulla A, Lewis CE, Carnethon MR, Schreiner PJ, et al. Gut microbiome and stages of diabetes in middle-aged adults: CARDIA microbiome study. Nutr Metab. 2023 Jan 5;20(1):3.

48. Chen Z, Radjabzadeh D, Chen L, Kurilshikov A, Kavousi M, Ahmadizar F, et al. Association of Insulin Resistance and Type 2 Diabetes With Gut Microbial Diversity: A Microbiome-Wide Analysis From Population Studies. JAMA Netw Open. 2021 Jul 1;4(7):e2118811.

49. Malaterre C, Dussault AC, Mermans E, Barker G, Beisner BE, Bouchard F, et al. Functional Diversity: An Epistemic Roadmap. BioScience. 2019 Oct 1;69(10):800–11.

50. Mouchet MA, Villéger S, Mason NWH, Mouillot D. Functional diversity measures: an overview of their redundancy and their ability to discriminate community assembly rules. Funct Ecol. 2010;24(4):867–76.

51. Qin J, Li Y, Cai Z, Li S, Zhu J, Zhang F, et al. A metagenome-wide association study of gut microbiota in type 2 diabetes. Nature. 2012 Oct;490(7418):55–60.

52. Winther SA, Mannerla MM, Frimodt-Møller M, Persson F, Hansen TW, Lehto M, et al. Faecal biomarkers in type 1 diabetes with and without diabetic nephropathy. Sci Rep. 2021 Jul 26;11(1):15208.

53. Chen J, Vitetta L. The Role of Butyrate in Attenuating Pathobiont-Induced Hyperinflammation. Immune Netw. 2020;20(2):e15.

54. Amiri P, Hosseini SA, Ghaffari S, Tutunchi H, Ghaffari S, Mosharkesh E, et al. Role of Butyrate, a Gut Microbiota Derived Metabolite, in Cardiovascular Diseases: A comprehensive narrative review. Front Pharmacol. 2022;12.

55. Tilg H, Moschen AR. Microbiota and diabetes: an evolving relationship. Gut. 2014 Sep 1;63(9):1513–21.

56. Li X, Zhang B, Hu Y, Zhao Y. New Insights Into Gut-Bacteria-Derived Indole and Its Derivatives in Intestinal and Liver Diseases. Front Pharmacol. 2021;12.

57. Ranhotra HS. Discrete interplay of gut microbiota L-tryptophan metabolites in host biology and disease. Mol Cell Biochem. 2023 Oct 20;

58. Zhang B, Jiang M, Zhao J, Song Y, Du W, Shi J. The Mechanism Underlying the Influence of Indole-3-Propionic Acid: A Relevance to Metabolic Disorders. Front Endocrinol. 2022;13.

59. Li CH, Wang CT, Lin YJ, Kuo HY, Wu JS, Hong TC, et al. Long-term consumption of the sugar substitute sorbitol alters gut microbiome and induces glucose intolerance in mice. Life Sci. 2022 Sep 15;305:120770.

60. Hossain KS, Amarasena S, Mayengbam S. B Vitamins and Their Roles in Gut Health. Microorganisms. 2022 Jun;10(6):1168.

61. Putnam EE, Goodman AL. B vitamin acquisition by gut commensal bacteria. PLOS Pathog. 2020 Jan 23;16(1):e1008208.

